# TRIP6 is required for tension at adherens junctions

**DOI:** 10.1101/2020.04.19.049569

**Authors:** Srividya Venkatramanan, Consuelo Ibar, Kenneth D. Irvine

## Abstract

Hippo signaling mediates influences of cytoskeletal tension on organ growth. TRIP6 and LIMD1 have each been identified as being required for tension-dependent inhibition of the Hippo pathway LATS kinases and their recruitment to adherens junctions, but the relationship between TRIP6 and LIMD1 was unknown. Using siRNA-mediated gene knockdown we show that TRIP6 is required for LIMD1 localization to adherens junctions, whereas LIMD1 is not required for TRIP6 localization. TRIP6, but not LIMD1, is also required for recruitment of Vinculin and VASP to adherens junctions. Knockdown of TRIP6 or Vinculin, but not of LIMD1, also influences the localization of phosphorylated myosin light chain and F-actin. In TRIP6 knockdown cells actin stress fibers are lost apically but increased basally, and there is a corresponding increase in recruitment of Vinculin and VASP to basal focal adhesions. These observations identify a role for TRIP6 in organizing F-actin and maintaining tension at adherens junctions that could account for its influence on LIMD1 and LATS. They also suggest that focal adhesions and adherens junctions compete for key proteins needed to maintain attachments to contractile F-actin.

## INTRODUCTION

The Hippo signaling network controls organ growth and cell fate in a wide range of animals, and when dysregulated, can contribute to oncogenesis (Misra and Irvine, 2018; Zanconato et al., 2016). Hippo signaling mediates its effects through regulation of the transcriptional co-activator proteins YAP1 and TAZ (Yorkie in *Drosophila*). YAP1 and TAZ (collectively, YAP proteins) are inhibited by Hippo signaling through phosphorylation by the LATS kinases LATS1 and LATS2 (Warts in *Drosophila*). Hippo signaling integrates diverse upstream inputs, including cytoskeletal tension. Distinct mechanisms through which cytoskeletal tension could modulate Hippo signaling have been suggested (Misra and Irvine, 2018; Sun and Irvine, 2016). One of the best characterized involves tension-dependent recruitment and inhibition of LATS kinases at adherens junctions (AJ). This was first discovered in *Drosophila*, where the Ajuba LIM protein (Jub) is recruited to AJ under tension, and Jub then recruits and inhibits Warts (Rauskolb et al., 2019; Rauskolb et al., 2014; Razzell et al., 2018). Tension-dependent inhibition and recruitment of LATS kinases to AJ has also been observed in mammalian cells, but two different proteins have been implicated in this recruitment: LIMD1 and TRIP6 (Dutta et al., 2018; Ibar et al., 2018).

Each of the three mammalian Ajuba family proteins: AJUBA, WTIP, and LIMD1, co-localize with LATS kinases at AJ, and have been reported to be able to co-immunoprecipitate with LATS kinases, and to inhibit them when over-expressed (Das Thakur et al., 2010; Ibar et al., 2018; Jagannathan et al., 2016). However, LIMD1 is the Ajuba family protein most closely related to *Drosophila* Jub, and is uniquely required in MCF10A cells for regulation of LATS at AJ under tension (Ibar et al., 2018). In the absence of LIMD1, or under conditions of low cytoskeletal tension, LATS kinases are not recruited to AJ, and LATS activity is increased. Localization of Ajuba family proteins to junctions requires α-catenin, and observations that Ajuba family proteins co-localize with and can co-precipitate α-catenin, together with identification of α-catenin mutations that constitutively recruit Jub, imply that they are recruited to junctions through interaction with α-catenin (Alégot et al., 2019; Ibar et al., 2018; Marie et al., 2003). As α-catenin undergoes a tension-dependent conformational change (Kim et al., 2015; Yao et al., 2014; Yonemura et al., 2010), conformation-dependent binding to α-catenin could explain tension-dependent recruitment of Ajuba proteins (Alégot et al., 2019; Ibar et al., 2018).

TRIP6 is a member of the Zyxin family of LIM domain proteins, which like the Ajuba family proteins have 3 LIM domains at their C-terminus (Yi and Beckerle, 1998). Zyxin, the best characterized family member, plays important roles in stabilizing and remodeling actin stress fibers subject to mechanical strain (Smith et al., 2014). A wide variety of roles for TRIP6 have been described, including effects on tumorigenesis, cell motility, telomere function, anti-apoptotic signaling, and transcription, and TRIP6 has been reported to interact with a wide variety of binding partners (Lin and Lin, 2011; Willier et al., 2011). Notably, in epithelial cell lines, TRIP6 can co-immunoprecipitate with LATS kinases, co-localize with LATS kinases at junctions, and is required for localization of LATS kinases to junctions under tension (Dutta et al., 2018). TRIP6 was recruited to junctions under tension in a Vinculin (VCL)-dependent process. As VCL localizes to AJ through association with a tension-induced “open” form of α-catenin (Kim et al., 2015; Yao et al., 2014; Yonemura et al., 2010), interaction with VCL could account for the tension-dependent recruitment of TRIP6 to AJ.

The observation that LIMD1 and TRIP6 both exhibit tension-dependent localization to AJ and are both required for recruitment and inhibition of LATS kinases at junctions, raised the question of how their effects on LATS kinases are related. By analyzing LIMD1 and TRIP6 localization in cells subject to siRNA-mediated knockdown of TRIP6 or LIMD1, respectively, we found that TRIP6 is required for recruitment of LIMD1 to junctions, but LIMD1 is not required for recruitment of TRIP6 to junctions. Analysis of markers of junctional tension revealed that TRIP6, but not LIMD1, is also required for normal tension at AJ, which could account for its role in recruiting LIMD1, while staining for F-actin revealed that apical F-actin organization is altered in the absence of TRIP6. Knockdown of TRIP6 also leads to an increase in basal F-actin stress fibers and an apparent re-localization of VCL and VASP from AJ to focal adhesions (FA), suggesting the existence of a competition between FA and AJ for proteins that stabilize tensile F-actin fibers, with TRIP6 promoting recruitment of these proteins to AJ.

## RESULTS

### TRIP6 is required for LIMD1 localization to AJ

LIMD1 and TRIP6 each exhibit tension-dependent localization to AJ, where they overlap with LATS proteins and with VCL (Dutta et al., 2018; Ibar et al., 2018). To directly compare their localization, MCF10A cells were cultured under low cell density conditions, which promotes tension at junctions. Fixed cells were then stained with antisera against both proteins. This revealed extensive co-localization of LIMD1 and TRIP6 at AJ, where both proteins also co-localize with E-cadherin (E-cad) and with LATS1 (Fig. 1).

**Fig 1.**
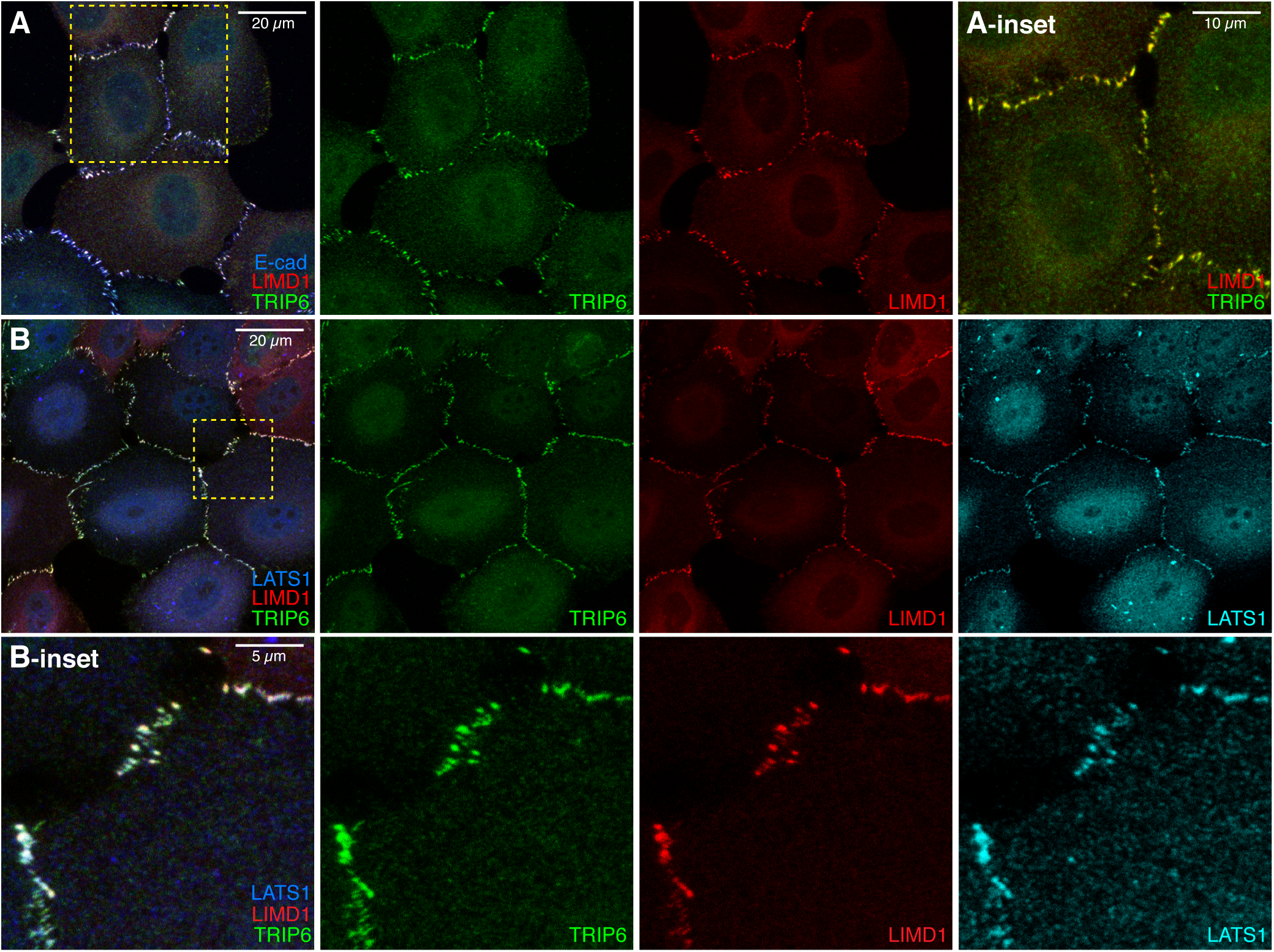
Co-localization of TRIP6, LIMD1 and LATS1 at AJ. (A-B) MCF10A cells plated at low density and cultured for 48 hours and then stained for mouse TRIP6 (green), LIMD1 (red) and E-cad (A) or LATS1 (blue/cyan) (B). Images are Z-projections through whole cells and are representatives of at least 3 biological replicates. Insets show higher magnification of the boxed regions, white bars indicate scale bars.

Since TRIP6 and LIMD1 co-localize at AJ, we next investigated whether localization of one protein depends upon the other. As TRIP6 siRNA was not completely effective in all cells, we used TRIP6 immunostaining to identify cells with strong knockdown of protein expression, and also examined knockdown using two independent siRNAs (Figs 2, S1). siRNA-mediated knockdown of TRIP6 substantially reduced LIMD1 and LATS1 localization to cell-cell junctions in cells cultured at low cell density, while E-cad staining remained similar to or only slightly weaker than that in control experiments (Figs 2A,B,M,N, S1A-F). Thus, TRIP6 is required for normal localization of LIMD1 at AJ. Conversely, siRNA-mediated knockdown of LIMD1 did not visibly reduce junctional localization of TRIP6, even though these same siRNA conditions clearly reduced LIMD1 levels (Fig. 2C,D). Moreover, functional knock-down of LIMD1 under these conditions was demonstrated by loss of the junctional localization of LATS1 (Fig. 2E,F), which depends upon LIMD1 (Ibar et al., 2018).

**Fig 2.**
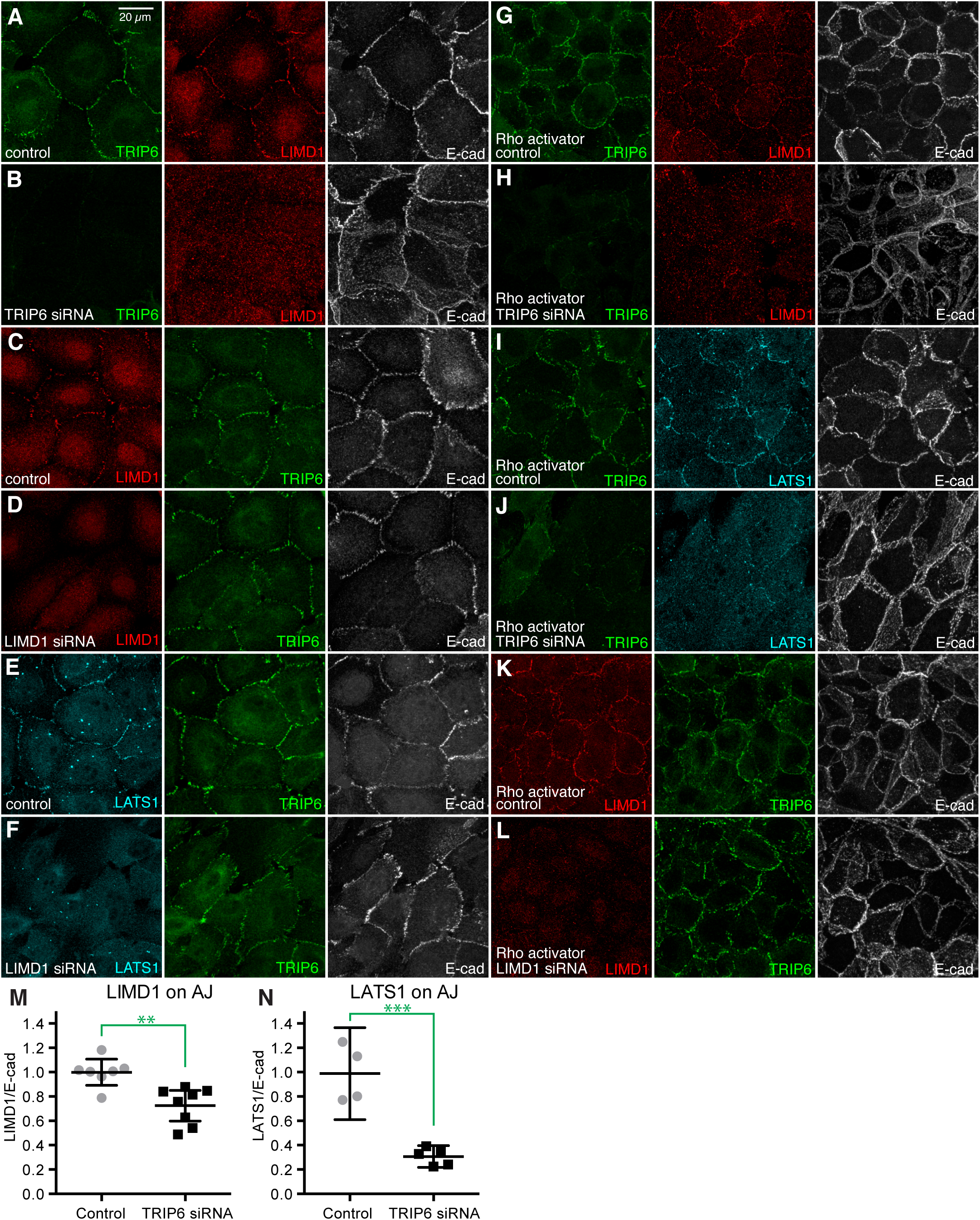
TRIP6 is required for junctional localization of LIMD1. (A-F) MCF10A cells plated at low density and transfected with control (A, C, E), TRIP6 (B) or LIMD1 (D, F) siRNA, cultured for 48 hours, then fixed and stained for TRIP6 (green), LIMD1 (red), LATS1 (cyan), or E-cad (white). (G-L) MCF10A cells plated at high density and transfected with control (G, I, K), TRIP6 (H, J) or LIMD1 (L) siRNA, and cultured for 48 hours and then treated with 1 µg/ml Rho-activator-II for 3 hours, fixed and stained for TRIP6 (green), LIMD1 (red) or LATS1 (cyan) or E-cad (white). Images are Z-projections through whole cells and are representatives of at least 3 biological replicates. (M, N) Quantification of junctional levels of LIMD1 and LATS1 normalized to junctional E-cad levels under both control and TRIP6 knockdown conditions. Each dot represents results from a confocal image stack containing several cells, as in the examples. For LIMD1 analysis N=7 for control, 8 for TRIP6 siRNA; for LATS1 analysis N=4 for control, 5 for TRIP6 siRNA. Significance of unpaired t tests indicated, ** indicates P<0.01, *** indicates P<0.001.

To further investigate the relationship between TRIP6 and LIMD1, we examined them under conditions where high cytoskeletal tension was established by treating cells with Rho activator II (Schmidt et al., 1997), rather than by culturing them at low cell density. Rho activator II increases F-actin and myosin activity, and thereby promotes localization of LIMD1 and LATS1 to AJ (Ibar et al., 2018). Junctional localization of LIMD1 and LATS1 was severely reduced by knockdown of TRIP6 even within Rho-activator treated cells (Fig. 2G-J, S1G-J). Conversely, knockdown of LIMD1 had no visible effect on junctional localization of TRIP6 in Rho activator treated cells (Fig. 2K,L). Altogether, these observations imply a hierarchical relationship between junctional localization of LIMD1 and TRIP6, in which TRIP6 is localized to AJ independently of LIMD1, whereas LIMD1 localization to AJ requires TRIP6. Moreover, they suggest that TRIP6 acts downstream or in parallel to the influence of Rho activation on LIMD1 and LATS1 localization.

### TRIP6 is required for cytoskeletal tension at AJ

The requirement for TRIP6 to localize LIMD1 could reflect a direct physical association that recruits LIMD1. Alternatively, since LIMD1 localization to AJ requires cytoskeletal tension, the requirement for TRIP6 might reflect an influence of TRIP6 on tension at AJ. To investigate this latter possibility, we examined VCL, as the localization of VCL to AJ depends upon a tension-induced conformational change in α-catenin (Kim et al., 2015; Yao et al., 2014; Yonemura et al., 2010). We observed that siRNA of TRIP6 diminished localization of VCL to AJ, whereas siRNA of LIMD1 had no noticeable effect (Fig. 3). This requirement for TRIP6 in VCL localization to AJ was observed both under low cell density conditions, and when cytoskeletal tension was elevated by treatment with Rho activator II. The finding that TRIP6 is required for junctional localization of VCL as well as of LIMD1 suggests that TRIP6 is required to maintain tension at AJ. We note that our observations differ from (Dutta et al., 2018), who reported that recruitment of VCL to junctions was not affected by knockdown of TRIP6, possibly due to differences in experimental conditions.

**Fig 3.**
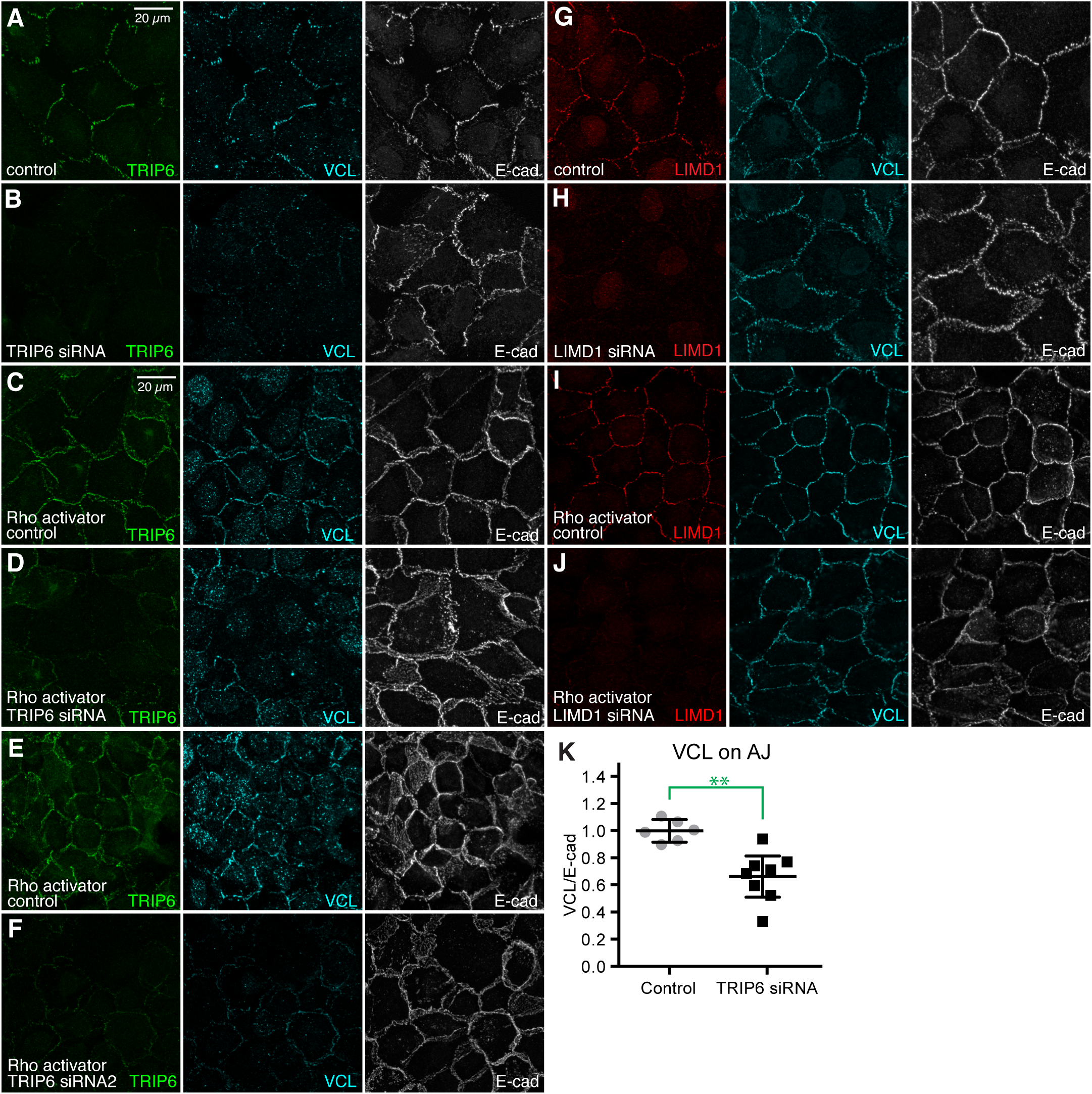
TRIP6 is required for VCL localization to AJ. (A-J) MCF10A cells plated at low (A, B, G, H) or high (C, D, E, F, I, J) density and transfected with control (A, C, E, G, I), TRIP6 (B, D, F) or LIMD1 (H, J) siRNA 24 hours after seeding. Cells grown at high density were then treated with 1 µg/ml Rho-activator-II for 3 hours. Cells were fixed and stained for TRIP6 (green), LIMD1 (red), VCL (cyan) anti-rabbit (A-D) or VCL anti-mouse (E-H), and E-cad (white). Images are Z-projections of confocal stacks and are representatives of at least 3 biological replicates. (K) Quantification of junctional levels of VCL normalized to junctional E-cad levels under control and TRIP6 knockdown conditions. Each dot represents results from a confocal image stack containing several cells, N=6 for control, 8 for TRIP6 siRNA. Significance of unpaired t test indicated, ** indicates P<0.01.

### TRIP6 influences the organization of actin and myosin

To investigate whether the apparent requirement for TRIP6 for tension at AJ reflects a more general influence on cytoskeletal organization, we examined the distribution of F-actin by staining with fluorescently-labelled phalloidin, and the distribution of active myosin using antibodies against phosphorylated myosin light chain (pMLC). Overall, we observed an apparent reduction in pMLC staining when TRIP6 was knocked down, and a reorganization of F-actin (Figs 4, S2). In contrast, knockdown of LIMD1 was not associated with any visible changes in staining for pMLC or F-actin (Fig. S2).

**Fig 4.**
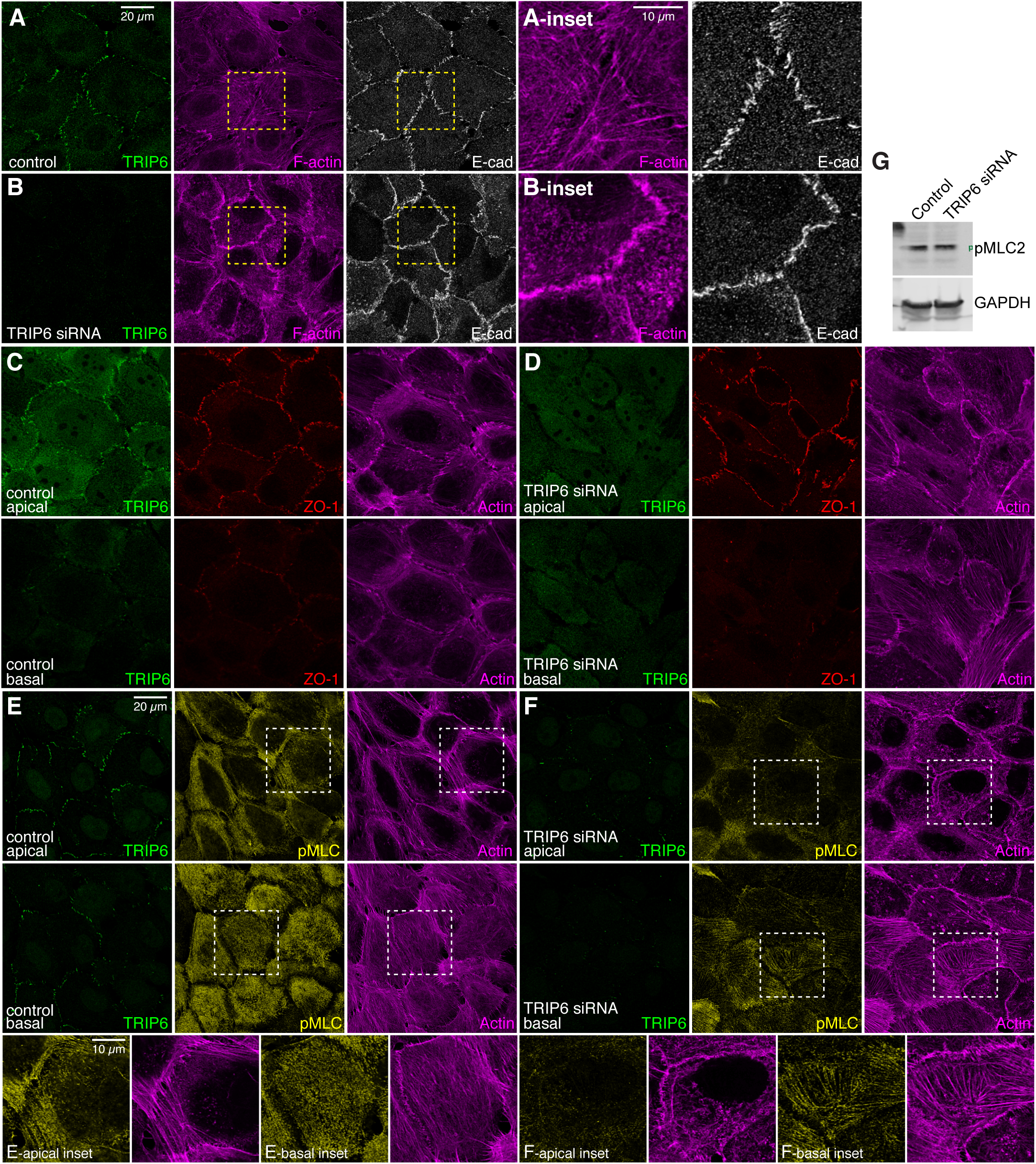
TRIP6 influences the organization of actin and myosin. (A-F) MCF10A cells plated at low density and transfected with either control or TRIP6 siRNA, as indicated, cultured for 72 hours and then fixed and stained for TRIP6 (green, A-B, E-F anti-mouse, C-D anti-rabbit), F-actin (magenta), and either E-cad (white) or ZO-1 (red) or pMLC (yellow). Images are projections through 3 to 7 Z-sections: apical sections (A-B) and (C-F: upper panels) or basal sections (C-F: lower panels) and are representatives of at least 3 biological replicates. Insets show higher magnification of the boxed regions. (G) Western blot showing pMLC2 and loading control (GAPDH) levels in control and TRIP6 siRNA cells.

In more apical regions (e.g. overlapping junctional staining for the tight junction protein ZO-1), MCF10A cells cultured at low density typically have multiple long F-actin fibers extending away from the AJ (Fig. 4A,C,E). In contrast, in TRIP6 knock-down MCF10A cells fewer of these long apical F-actin fibers are observed, and instead F-actin is often concentrated in thick, irregular accumulations along the AJ, as well as diffuse speckles of F-actin within the cytoplasm (Fig. 4B,D,F). This is accompanied by a reduction in apical pMLC staining (Figs 4E,F, S2A,C).

In more basal regions (e.g. basal to junctional staining for ZO-1), MCF10A cells often appear to have a meshwork of thin F-actin filaments (Fig. 4C,E). In TRIP6 knockdown MCF10A cells, basal F-actin is instead often observed to concentrate in thick F-actin fibers that run along the basal surface of the cell (Fig. 4D,F). This reorganization of F-actin is accompanied by altered pMLC staining, from diffuse basal puncta in wild-type MCF10A cells (Fig. 4E), to concentration along basal F-actin fibers in TRIP6 knock-down cells (Fig. 4F). Despite the overall appearance of reduced pMLC immunostaining, no reduction in pMLC levels was detected by western blotting with anti-pMLC antibodies (Fig. 4G), implying that TRIP6 siRNA leads to a change in pMLC localization, rather than a decrease in total pMLC levels within the cell.

To quantify these differences in F-actin, we used a published MATLAB script for segmentation and analysis of actin fibers (Rogge et al., 2017). This confirmed the visual impression of a reduction in apical actin fibers, and an increase in basal actin fibers (Fig 5A-E). It also revealed a relative decrease in the length of apical actin fibers, and an increase in the width of basal actin fibers (Fig. 5F,G). These observations indicate that knockdown of TRIP6 is not simply associated with a reduction in cytoskeletal tension. Rather, there is a reorganization of cytoskeletal tension that includes an apparent reduction in apical actin stress fibers attached to AJ, and an increase in basal actin stress fibers.

**Fig. 5.**
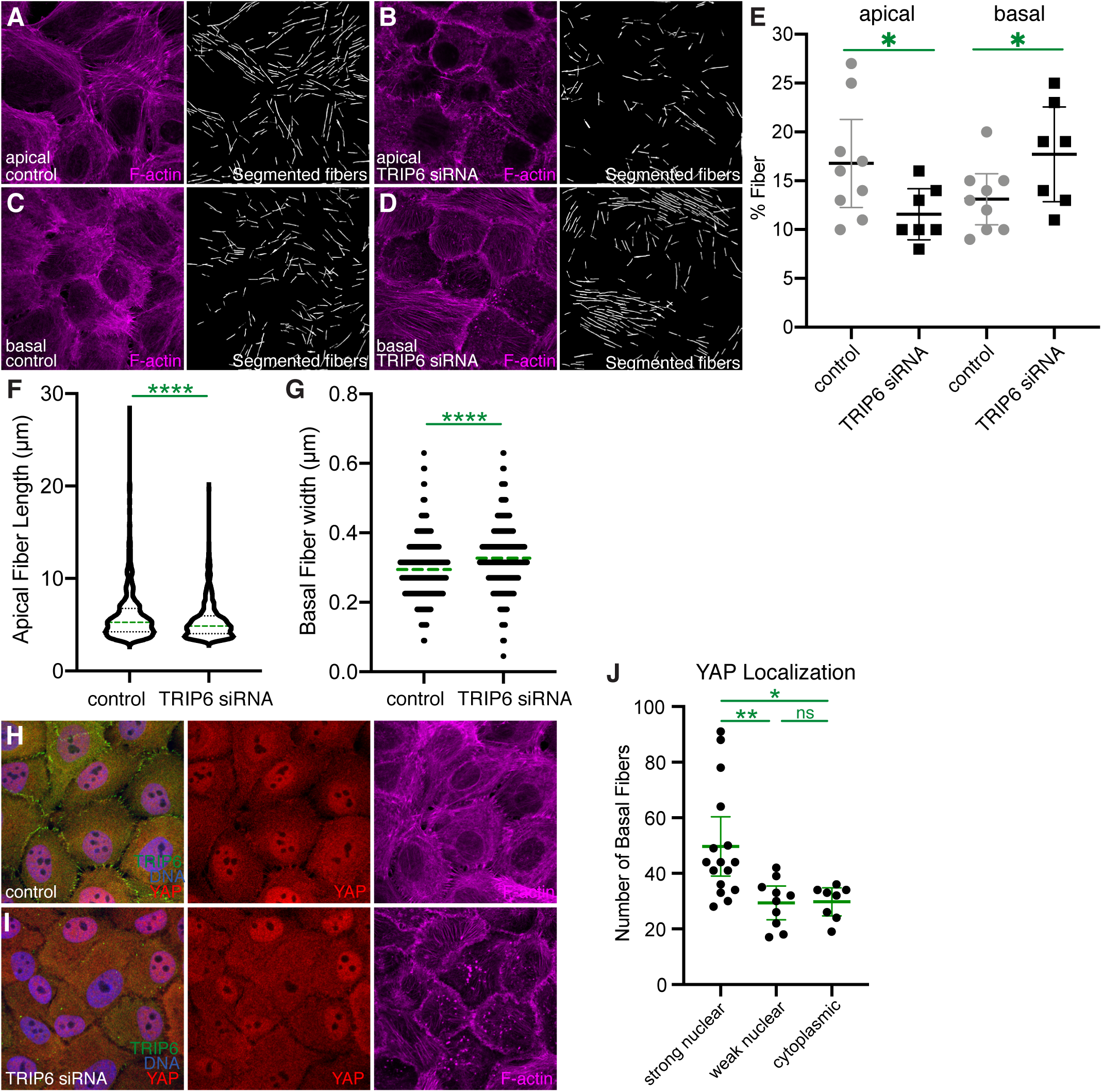
Quantitation of actin fibers in TRIP6 knockdown cells. (A-D) Examples of F-actin segmentation using FSegment, in MCF10A cells plated at low density and transfected with either control or TRIP6 siRNA. F-actin staining from apical or basal regions, together with segmented actin fibers, are shown. (E) Comparison of percent of the image occupied by F-actin fibers in image panels similar to those shown in A-D. Each dot represents one confocal image, comprising several cells as in the examples. Bars indicate mean and 95% confidence interval. (F) Comparison of the length distribution of apical actin fibers in control versus TRIP6 RNAi images, displayed in a violin plot with the median indicated by a dashed green line. N=816 fibers in control, 496 fibers in TRIP6 RNAi. (G) Comparison of the width distribution of basal actin fibers in control versus TRIP6 RNAi images, displayed in a dot plot with the median indicated by a dashed green line. Steps between points reflect 1 pixel differences in width. N=788 fibers in control, 709 fibers in TRIP6 RNAi. (H, I) Examples of YAP (red) localization and F-actin (magenta) staining in MCF10A cells plated at low density and transfected with either control or TRIP6 siRNA. (J) Comparison of the number of basal actin fibers in TRIP6 RNAi treated cells with strong nuclear YAP, weak nuclear YAP, or no elevation in nuclear YAP above cytoplasmic levels, displayed in a dot plot with the mean and 95% CI indicated by green bars. Each dot represents one cell. Results of statistical comparisons performed by t test in E, F and G, and by one-way ANOVA in J are indicated in green, * indicates P<0.05, ** indicates P<0.01, *** indicates P<0.001, **** indicates P<0.0001, ns indicates not significant.

Knockdown of TRIP6 was previously reported to be associated with reduced YAP activity (Dutta et al., 2018). As many studies have linked F-actin levels and tension in the actin cytoskeleton to YAP activation (Misra and Irvine, 2018; Sun and Irvine, 2016), we wondered whether TRIP6 knockdown could reduce YAP activity even under conditions where prominent basal actin fibers were observed. Under our low cell density culture conditions, YAP is predominantly nuclear in MCF10A cells (Fig 5H). The overall YAP distribution was not consistently affected by TRIP6 knockdown in our experiments (Fig 5I). However, there was cell-to-cell variation both in the extent of formation of basal F-actin stress fibers in TRIP6 knock-down cells, and in the YAP localization profile. When we quantified the number of basal F-actin fibers in individual TRIP6 knock down cells, and compared this to the YAP localization profile, we observed a correlation between nuclear YAP localization and higher numbers of basal F-actin fibers as compared to cells without reduced nuclear YAP localization (Fig. 5J). These observations suggest that the expected decrease in YAP activity generated by loss of TRIP6 from apical junctions might be compensated for by an increase in YAP activity generated by elevated basal actin stress fibers.

### Re-localization of VCL and VASP in the absence of TRIP6

To explain the shift in F-actin stress fibers from apical to basal in TRIP6 knockdown cells, we hypothesized that there could be a competition between apical AJ and basal FA for protein(s) that are required for maintenance or attachment of F-actin stress fibers, and present in limiting amounts within cells. According to this model TRIP6 could recruit these proteins to AJ, but in the absence of TRIP6 they would instead be recruited to FA (Fig. 6D). To begin to investigate this possibility, we examined the distribution of proteins that can localize to AJ and FA, and can influence or interact with F-actin. One such protein is VCL. Confluent MCF10A cells lack prominent FA, and basal accumulation of VCL is difficult to detect (Fig. 6B). However, in TRIP6 knockdown cells, basal puncta of VCL are readily apparent, particularly at the ends of F-actin stress fibers (Fig. 6C). These observations imply that knockdown of TRIP6 promotes formation of VCL-containing FA.

**Fig 6.**
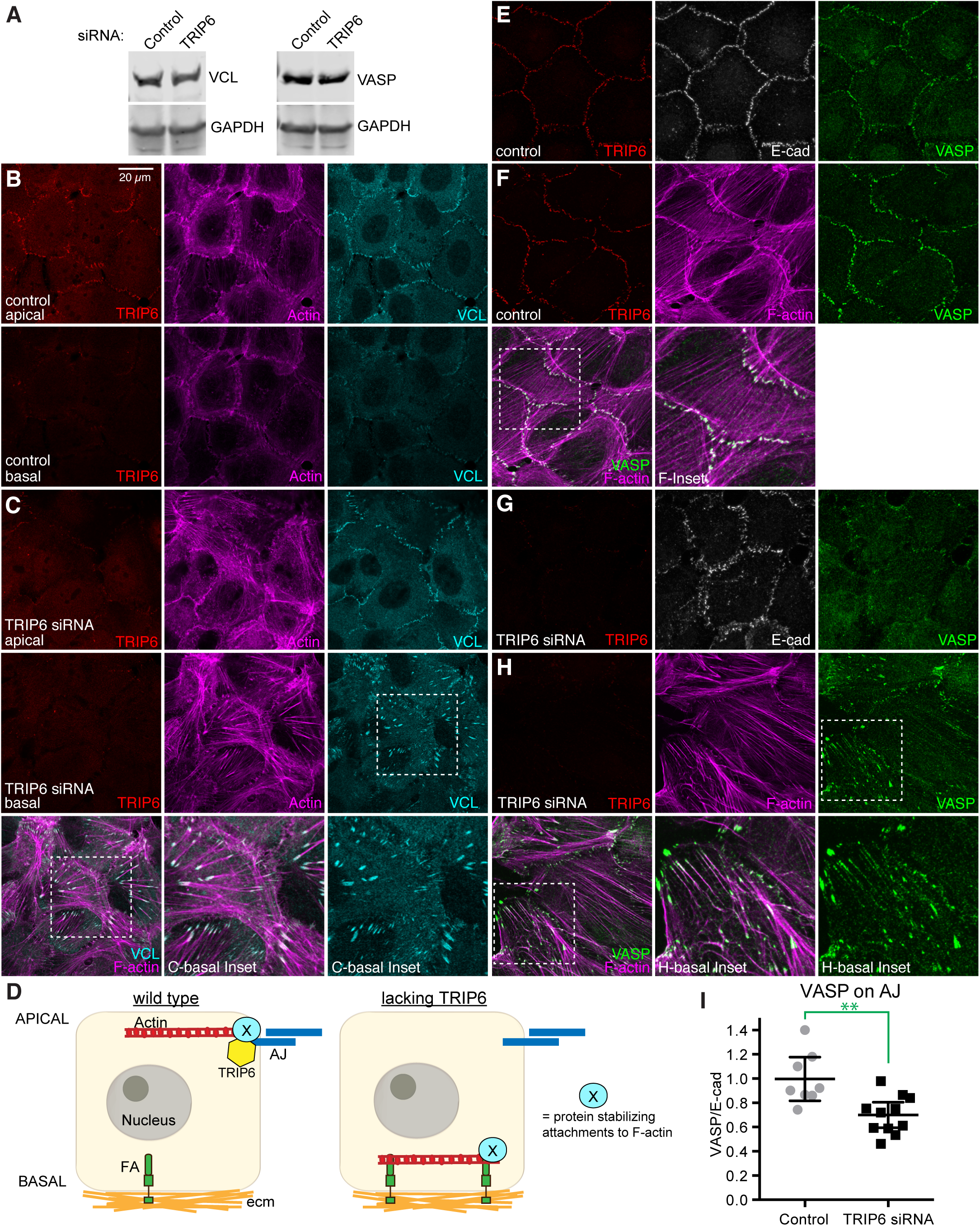
TRIP6 knockdown shifts localization of VCL and VASP to focal adhesions. (A) Western blots showing VCL or VASP and loading control (GAPDH) levels in control and TRIP6 siRNA cells. (B-C, E-H) MCF10A cells plated at low-density and transfected with either control or TRIP6 siRNA, cultured for 72 hours and then fixed and stained for TRIP6 (red) (anti-rabbit A-B or anti-mouse C-F), E-cad (white) or F-actin (magenta) and VCL (cyan) (anti-mouse) or VASP (green). Images are projections through either whole cell (E-G) or 3 to 7 apical (upper panel) and basal (lower panel) sections and are representatives of at least 3 biological replicates. Insets show higher magnification of the boxed regions. (D) Model illustrating influence of TRIP6 on actin stress fibers. In presence of TRIP6 (left), TRIP6 recruits a crucial actin regulatory protein (X), which stabilizes attachments of tensile F-actin. In the absence of TRIP6, X is instead recruited to FA, where it stabilizes basal F-actin stress fibers. (I) Quantification of junctional levels of VASP normalized to junctional E-cad levels under control and TRIP6 knockdown conditions. Each dot represents results from a confocal image stack containing several cells, N=8 for control, 11 for TRIP6 siRNA. Significance of unpaired t test indicated, ** indicates P<0.01.

VASP is an actin regulatory protein that can be observed at AJ in epithelial cells, but at FA and leading edge in fibroblasts (Krause et al., 2003). Prominent VASP localization to AJ was observed in wild-type MCF10A cells (Fig. 6E,F). When TRIP6 is knocked down, VASP is lost from AJ, and is instead detected in basal puncta (Fig. 6G-I). Western blotting did not reveal any change in the total levels of VCL or VASP in TRIP6 knockdown cells (Fig. 6A). The basal puncta of VASP are most prominent at the ends of F-actin stress fibers, presumably representing accumulation at FA, but some VASP accumulation can also be observed along basal stress fibers (Fig. 6H). Thus, both VCL and VASP re-localize from AJ to FA in TRIP6 knockdown cells, coincident with the reorganization of F-actin and pMLC.

### VCL is required for tension at adherens junctions and focal adhesions

While our observations show that TRIP6 is required for normal localization of VCL to AJ, published observations have reported that TRIP6 localization to AJ is dependent upon VCL (Dutta et al., 2018). To further investigate the relationship between VCL and TRIP6, and their influence on the cytoskeleton and Hippo signaling, we compared the consequences of VCL knockdown on tension-regulated proteins at AJ and the actomyosin cytoskeleton in MCF10A cells.

VCL siRNA reduced junctional localization of LIMD1 and resulted in a substantial reduction in pMLC staining, similar to TRIP6 knockdown (Fig. 7A-F). However, in other respects, the consequences of VCL knockdown differed. Strong VCL knockdown consistently reduced junctional localization of E-cad (Fig. 7A-C), whereas TRIP6 knockdown had much less effect on E-cad. VCL knockdown reduced apical F-actin stress fibers attached to AJ similar to TRIP6 knockdown (Fig. 7D-I), but did not lead to the appearance of basal F-actin stress fibers (Fig. 7H, I). Moreover, instead of thick F-actin accumulations along AJ, in apical regions VCL knockdown often resulted in the appearance of F-actin spikes (Fig. 7E, F, I). These phenotypic differences emphasize that TRIP6 and VCL have distinct roles at AJ.

**Fig 7.**
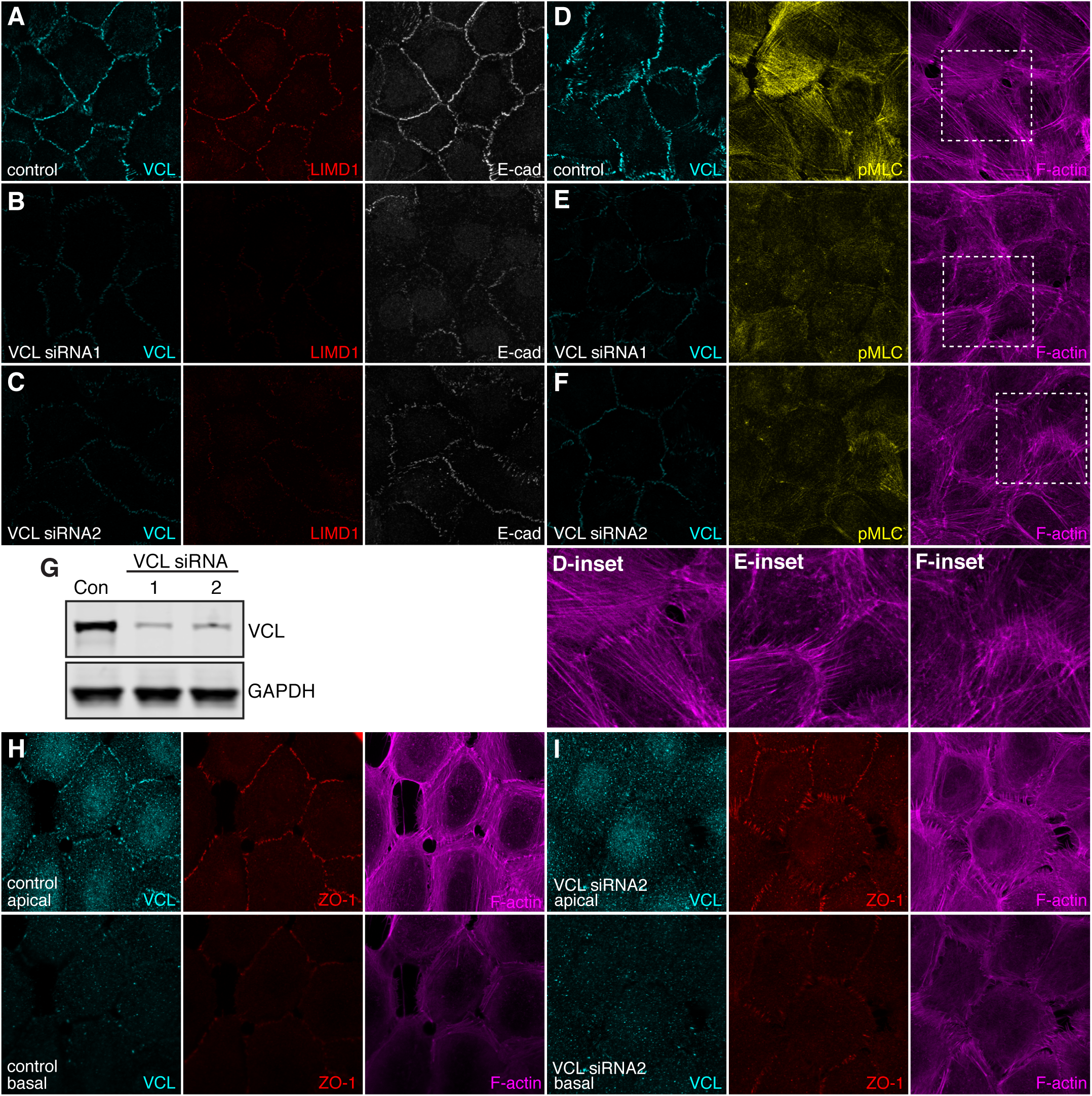
VCL is required to maintain tension at AJ. (A-F) MCF10A cells plated at low-density and transfected twice with either control or VCL siRNA, cultured for 48 hours then fixed in the presence of 0.65% Triton-X-100 and stained with mouse VCL (cyan), E-cad (white) or Phalloidin (magenta), rabbit LIMD1 (red) or rabbit pMLC (yellow). Images are Z-projections of confocal stacks and are representatives of at least 3 biological replicates. (G) Western Blot showing the knockdown efficiency of the VCL siRNAs used. Con, control (negative control siRNA). GAPDH is the loading control for the blot. (H-I) MCF10A cells plated at low density, transfected with either control or VCL siRNA and stained for rabbit VCL (cyan), mouse ZO-1 (red) and F-actin (magenta). ZO-1 staining was used to separate the images into apical (top panel) and basal sections (bottom panel).

## DISCUSSION

Prior studies ascribed similar effects on Hippo signaling to two LIM domain proteins, LIMD1 and TRIP6 (Dutta et al., 2018; Ibar et al., 2018). Our studies have now established distinct activities for these proteins, as we find that TRIP6 is required for localization of LIMD1 to AJ, whereas LIMD1 does not affect localization of TRIP6. The influence of TRIP6 on LIMD1 could in principle account its reported effects on Hippo signaling, though we note our results do not exclude the possibility that TRIP6 also influences Hippo signaling through other mechanisms (Dutta et al., 2018). Our results have further identified requirements for TRIP6 in localization of VCL and VASP, and in organization of F-actin.

Both LIMD1 and VCL are recruited to an open form of α-catenin that is generated under tension (Alégot et al., 2019; Ibar et al., 2018; Kim et al., 2015; Yao et al., 2014; Yonemura et al., 2010). The requirement for TRIP6 in localization of LIMD1 and VCL to AJ thus implies that it is required to maintain this open form. Consistent with this, we observe that knockdown of TRIP6 results in loss of pMLC-associated F-actin stress fibers attached to AJ. These observations thus suggest that TRIP6 is required to establish or maintain attachments of tensile actin stress fibers to AJ.

Loss of TRIP6 is not only associated with a loss of apical actin stress fibers attached to AJ, but also with a gain of basal actin stress fibers attached to FA. Intriguingly, loss of E-cad in MCF10A cells has also been reported to result in the formation of thick basal actin stress fibers (Chen et al., 2014). Although it was reported that apical F-actin did not change in E-cad mutant MCF10A cells, in those experiments wild-type MCF10A cells did not exhibit the tensile apical F-actin that we observe, possibly due to higher cell density or other aspects of culture conditions. To explain the reorganization of F-actin that occurs when TRIP6 is knocked down, we suggest that there could be a competition between AJ and FA for a limiting pool of proteins needed to stabilize attachments to actin stress fibers, and that TRIP6 is needed to recruit these proteins to AJ. When TRIP6 is eliminated, these proteins are then instead able to associate with FA, where they promote and stabilize basal stress fibers. We identified two proteins that are candidates to be affected by this hypothesized competition, VCL and VASP. Their localization shifts from predominantly at AJ in wild-type MCF10A cells, to predominantly at FA in TRIP6 knockdown MCF10A cells.

VCL is recruited through tension-sensitive mechanisms to AJ and FA, where it binds to F-actin, helps to stabilize association of F-actin to AJ and FA, and plays crucial roles in maintaining cell-cell and cell-matrix adhesions (Bays and DeMali, 2017). Although both VCL and TRIP6 knockdown resulted in loss of apical F-actin stress fibers, in other respects their influence on actin organization differed. VCL knockdown leads to loss rather than gain of basal stress fibers. Apically, VCL knockdown led to F-actin spikes forming along cell-cell junctions. As there was a noticeable reduction of E-cad at junctions in VCL knockdown cells, the formation of F-actin spikes might be related to a recently described process in which apical actin protrusions are suggested to play a role in repair of failing adhesive junctions (Li et al., 2020).

VASP regulates actin polymerization, and VCL and VASP have been reported to be able to interact with each other, and to function together in promoting tension-sensitive actin polymerization at AJ (Leerberg et al., 2014). VASP has also been found to be able to associate with TRIP6 and other related LIM domain proteins, including Zyxin and LPP, supporting the possibility of a direct role for TRIP6 in recruiting VASP (Hoffman et al., 2006; Petit et al., 2003; Reinhard et al., 1995). Alternatively, the influence of TRIP6 on VCL and VASP recruitment could be indirect. Indeed, as these proteins are recruited to sites of tension at AJ and FA, and they help to maintain tension at these same sites, they participate in a positive feedback loop that maintains attachments to actin stress fibers and thus their own recruitment to these sites.

Although our results indicate that TRIP6 plays an essential role in attachments of stress fibers to AJ, it was not required at FA. Studies in other cell types have yielded conflicting results on the effects of TRIP6 on FA (Lin and Lin, 2011; Willier et al., 2011). For example, in fibroblasts over-expressed TRIP6 was reported to bind to Supervillin to suppress FA maturation (Takizawa et al., 2006), and in A509 cells TRIP6 knockdown increased basal stress fibers (Guryanova et al., 2005). Conversely, in Hela cells TRIP6 was reported to facilitate maturation of FAs (Bai et al., 2007), and knockdown of TRIP6 in human endothelial cells was reported to decrease formation of actin fibers (Sanz-Rodriguez et al., 2004). In addition to TRIP6, the related Zyxin family proteins LPP and Zyxin have also been implicated in actin stress fiber organization (Smith et al., 2014), and they might contribute to stabilization of basal stress fibers in the absence of TRIP6.

Knockdown of TRIP6 was previously reported to lead to reduced nuclear YAP localization and reduced YAP activity (Dutta et al., 2018). We did not observe an overall reduction in nuclear YAP in our experiments, but we note that Dutta et al. (2018) did not report the increase in basal actin stress fibers that we observed. Thus, the distinct effects on YAP observed might reflect differences between experimental conditions that result in different consequences for basal F-actin. Several mechanisms by which basal actin stress fibers could contribute to nuclear YAP localization have been described (Misra and Irvine, 2018), including effects on integrin signaling and on nuclear pore size (Elosegui-Artola et al., 2017; Kim and Gumbiner, 2015). The correlation between basal F-actin and nuclear YAP in TRIP6 knockdown cells suggests that loss of TRIP6 switches the predominate mechanism for cytoskeletal regulation of Hippo signaling in MCF10A cells from dependence on LATS regulation at AJ to mechanisms that depend upon FA and basal F-actin stress fibers.

## MATERIALS AND METHODS

### Plasmids and siRNAs

Sequences of the siRNAs (5’-> 3’) used were as follows: human TRIP6 siRNA1-CCAAUGUUCCACUUUUGGUAUUGAT (IDT), human TRIP6 siRNA2-GCUCUGGAUCGAAGUUUUCACAUTG (IDT), human VCL siRNA1-CCAGUGGAUCGAUAAUGUUGAAAAA (IDT), human VCL siRNA2-GGCAAAUCAGUUACUAAGAAGAAAA (IDT). siRNAs used for LIMD1 have been described previously (Ibar et al., 2018).

### Cell culture

MCF10A cells (a gift from Jay Debnath, UCSF, CA) were cultured in DMEM/F12 supplemented with 5% horse serum, epidermal growth factor (20 µg/ml), insulin (10 µg/ml), cholera toxin (0.1 µg/ml), hydrocortisone (0.5 µg/ml) and antibiotic-antimycotic at 37°C along with 5% CO_2_. Cells were used at low passage number and checked for contamination by cell morphology and mycoplasma testing. Coverslips were coated with 0.6 mg/ml of collagen for 15 minutes at room temperature and washed with PBS before plating cells. For low cell density experiments, cells were grown at densities of around 35,000 cells/cm^2^. For Rho-activator experiments, cells were grown at high density (150,000 cells/cm^2^) for 48 hours and were then treated with Rho-activator-II (Cytoskeleton: 1 µg/ml) for 3 hours. For siRNA experiments, Lipofectamine RNAiMax (Life Technologies) was used to deliver siRNA into MCF10A cells following manufacturer’s protocol and cells were fixed and stained 72 hours after transfection (48 hours for LIMD1 siRNA transfection).

### Immunostaining and Imaging

Cells were either fixed with 4% paraformaldehyde in PBS++ (phosphate-buffered saline supplemented with 100 mM MgCl_2_ and 50 mM CaCl_2_) for 10 min at room temperature, or, when staining for VCL, LIMD1, or VASP, fixed in 1% paraformaldehyde with 0.65% Triton X-100 in PBS++ for 3 min, rinsed and then fixed in 1% PFA in PBS++ for 10 min. The cells were then washed three times 10 mins each with 200 mM glycine containing PBS, followed by permeabilization with 0.5% Triton X-100 in PBS for 20 min. After blocking with 5% bovine serum albumin (BSA) in PBS for 1 h, cells were incubated with primary antibody diluted in a 5% BSA in PBS solution overnight at 4°C. (pMLC staining-done over weekend). After washing with PBS, cells were incubated with Alexa Fluor 488-(Life Technologies), Cy3- or Alexa Fluor 647-conjugated (Jackson ImmunoResearch) secondary antibodies for 1 h and washed four times with PBS. Cell nuclei were counterstained with Hoechst 33342 (1 µg/ml; Invitrogen) and mounted with mounting medium (Dako). Antibodies used for immunostaining include mouse anti-Yap (1:100; Santa Cruz Biotechnology, sc-101199), rabbit anti-LATS1 (1:600; Cell Signaling Technology, #3477), mouse anti-VCL (1:2000; Sigma, V9131), rabbit anti-VCL (1:100; EMD Millipore, #MAB3574), rabbit anti-VASP (1:100; Cell Signaling Technology #3132S), mouse anti-ZO-1 (1:1000; Life Technology, #33-9100), mouse anti-pMLC (S19) (1:200; Cell Signaling Technology, #3675), rabbit anti-pMLC (T18/S19) (1:50; Cell Signaling Technology, #3674), rat anti-E-cadherin (1:500; Life Technology, #13-1900), rabbit anti-LIMD1 (1:500; Bethyl, A303-182A), rabbit anti-TRIP6 (1:50; Abcam, ab70747) and mouse anti-TRIP6 (1:100; Santa Cruz Biotechnology sc-365122). F-actin was stained with Alexa Fluor 647-conjugated phalloidin (1:50; Life Technologies). Images were acquired using LAS X software on a Leica TCS SP8 confocal microscope system using a HC PL APO 63×/1.40 objective.

### Actin segmentation and quantitation

Actin stress fibers were quantified using the FSegment MATLAB script developed by Rogge et al. (2017). This script uses a trace algorithm to trace linear structures of actin and suppresses non-linear fluorescence. Segmented linear structures are then analyzed with respect to fluorescence values for the actin fibers. The application allows calculation of fiber width, fiber length, total number of actin filaments, proportion of actin in the form of stress fibers compared to total actin (% Fiber actin), and total filament length. For YAP localization vs. Basal F-actin correlation studies in the TRIP6 siRNA treated cells, individual cells were cropped using the ‘FreeHandCrop’ feature in the Preprocessing GUI and the actin SFs analysis for each cell was then compared to the YAP localization within the cell. For comparison of actin between control and TRIP6 siRNA treated cells in the apical and basal sections, the entire image field was segmented and analyzed. Individual z-section representative of the apical and basal sections of the whole image field were selected for these analyses.

### Immunoblotting

MCF10A cells were lysed in 2x Laemmli sample buffer (Bio-rad) supplemented with protease inhibitor cocktail (Roche) and phosphatase inhibitor cocktail (Calbiochem). Protein samples were loaded to 4%-15% gradient gels (Bio-rad). Antibodies for immunoblotting include rabbit anti-vinculin (1:5000; EMD Millipore, #MAB3574), mouse anti-phospho-myosin light chain (S19) (1:500; Cell Signaling Technology, #3675), mouse anti-TRIP6 (1:1000; Santa Cruz Biotechnology sc-365122), rabbit anti-VASP (1:1000; Cell Signaling Technology #3132S) and rabbit-myosin light chain (1:1000; Cell Signaling Technology #3672S). As loading control, rabbit anti-GAPDH (1: 10,000; Santa Cruz Biotechnology sc-25778) or mouse anti-GAPDH (1:10,000; Novus Biologicals, NBP2-27103) were used. Blots were stained with Li-COR fluorescent-conjugated secondary antibodies at 1:20,000 dilution and were visualized and quantified using Odyssey Imaging System (Li-COR Biosciences).

### Statistical Analysis

Statistical comparisons were performed using Excel and GraphPad Prism software. For ratio comparisons, analysis was done on the log transform of the ratio.

## ACKNOWLEDGMENTS

We thank Jay Debnath for MCF10A cells, and Dan McCollum for helpful discussions. This research was supported by National Institutes of Health grant GM131748 (KDI) and a Busch pre-doctoral fellowship (SV)

## Supplemental Figures

**Fig. S1.**
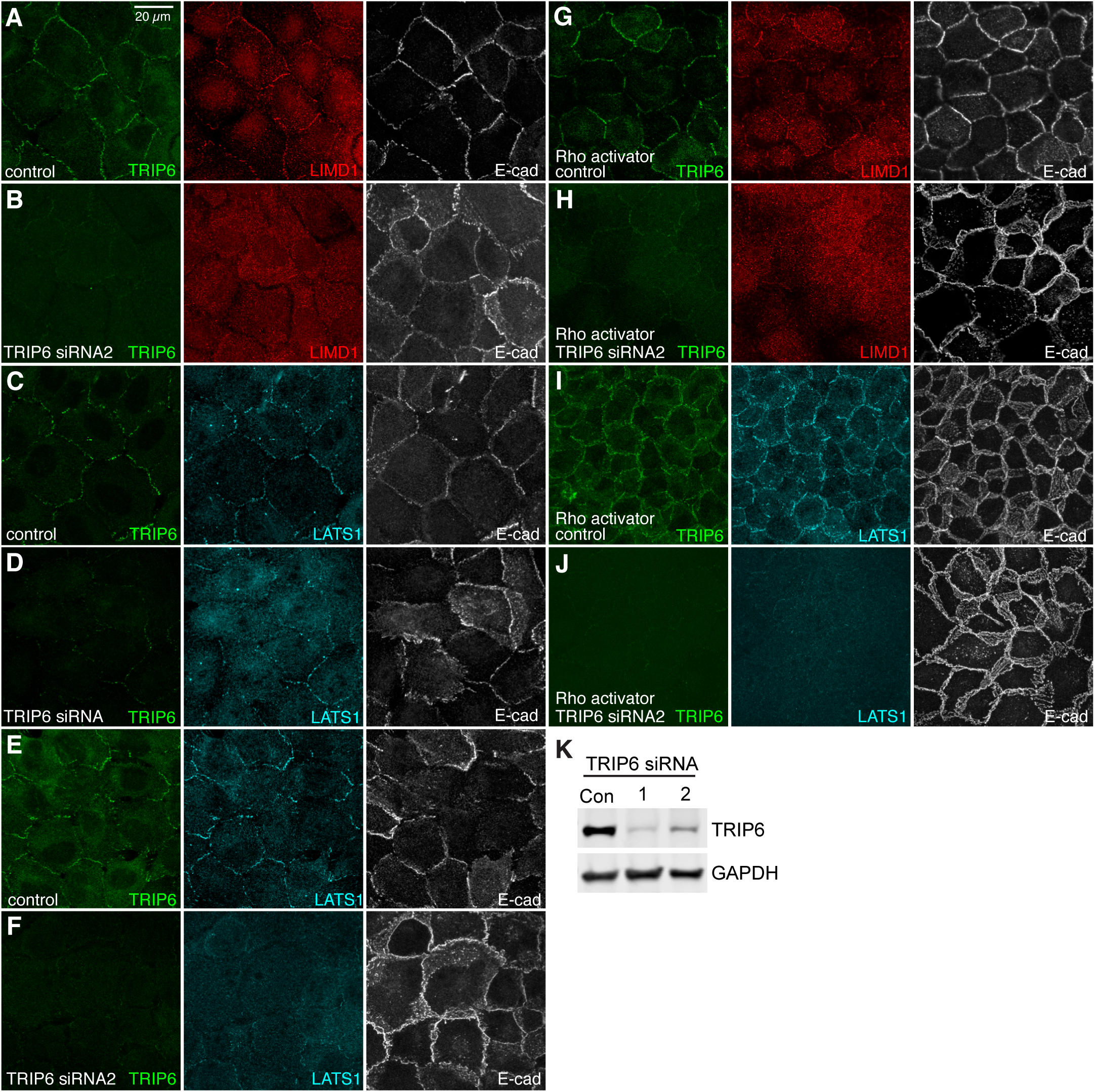
Validation of TRIP6 siRNAs and their effect on LIMD1 and LATS1 localization. (A-J) MCF10A cells grown at low density and transfected with control or TRIP6 siRNA or siRNA2 as indicated, cultured for 72 hours, then fixed and stained for TRIP6 (green) (anti-mouse), LIMD1 (red) or LATS1 (cyan), and E-cad (white). MCF10A cells in (G-J) were cultured at high cell density and also treated with 1 µg/ml Rho-activator-II for 3 hours before fixation. (K) Western Blot indicating knockdown efficiency of TRIP6 siRNAs used. Con, control (negative control siRNA). GAPDH is the loading control.

**Fig. S2.**
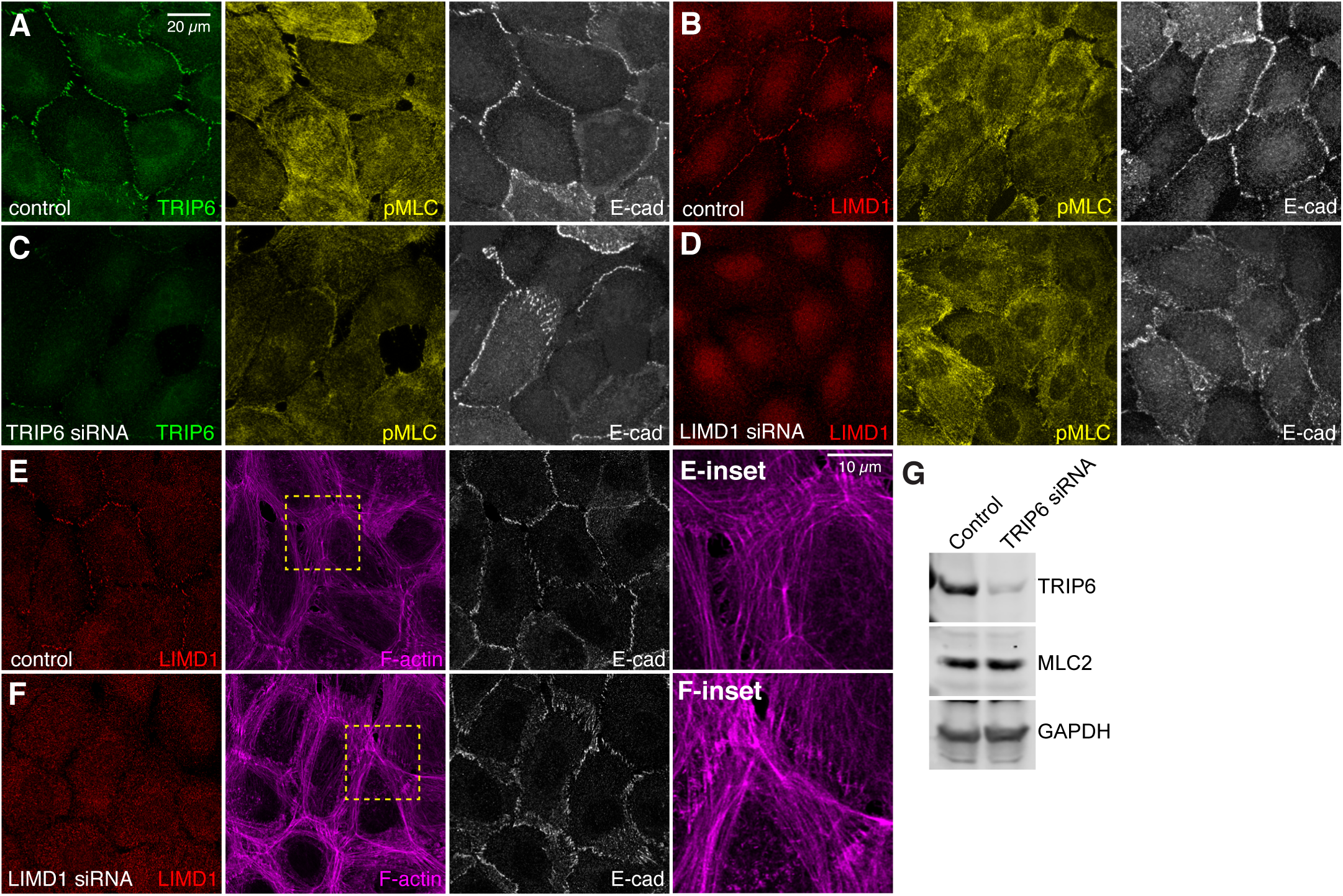
Effect of TRIP6 or LIMD1 knockdown on F-actin and pMLC. (A-F) MCF10A cells at low density and transfected with control, TRIP6 or LIMD1 siRNA, cultured for 48 hours (LIMD1 siRNA) or 72 hours (TRIP6 siRNA), then fixed and stained for TRIP6 (green) (anti-mouse) or LIMD1 (red), pMLC (yellow) or Phalloidin (magenta) and E-cad (white). (G) Western blot showing pMLC2 and loading control (GAPDH) levels in control and TRIP6 siRNA cells.

## REFERENCES

Alégot, H., C. Markosian, C. Rauskolb, J. Yang, E. Kirichenko, Y.-C. Wang, and K.D. Irvine. 2019. Recruitment of Jub by alpha-catenin promotes Yki activity and Drosophila wing growth. J Cell Science. 132:jcs222018.

Bai, C.-Y., M. Ohsugi, Y. Abe, and T. Yamamoto. 2007. ZRP-1 controls Rho GTPase-mediated actin reorganization by localizing at cell-matrix and cell-cell adhesions. Journal of Cell Science. 120:2828–2837.

Bays, J.L., and K.A. DeMali. 2017. Vinculin in cell–cell and cell–matrix adhesions. Cellular and molecular life sciences: CMLS. 74:2999–3009.

Chen, A., H. Beetham, M.A. Black, R. Priya, B.J. Telford, J. Guest, G.A.R. Wiggins, T.D. Godwin, A.S. Yap, and P.J. Guilford. 2014. E-cadherin loss alters cytoskeletal organization and adhesion in non-malignant breast cells but is insufficient to induce an epithelial-mesenchymal transition. BMC cancer. 14:3756–3714.

Das Thakur, M., Y. Feng, R. Jagannathan, M.J. Seppa, J.B. Skeath, and G.D. Longmore. 2010. Ajuba LIM proteins are negative regulators of the Hippo signaling pathway. Curr Biol. 20:657–662.

Dutta, S., S. Mana-Capelli, M. Paramasivam, I. Dasgupta, H. Cirka, K. Billiar, and D. McCollum. 2018. TRIP6 inhibits Hippo signaling in response to tension at adherens junctions. EMBO reports. 19:337–350.

Elosegui-Artola, A., I. Andreu, A.E.M. Beedle, A. Lezamiz, M. Uroz, A.J. Kosmalska, R. Oria, J.Z. Kechagia, P. Rico-Lastres, A.-L. Le Roux, C.M. Shanahan, X. Trepat, D. Navajas, S. Garcia-Manyes, and P. Roca-Cusachs. 2017. Force Triggers YAP Nuclear Entry by Regulating Transport across Nuclear Pores. Cell. 171:1397–1410.e1314.

Guryanova, O.A., A.A. Sablina, P.M. Chumakov, and E.I. Frolova. 2005. Downregulation of TRIP6 Gene Expression Induces Actin Cytoskeleton Rearrangements in Human Carcinoma Cell Lines. Molecular Biology. 39:792–795.

Hoffman, L.M., C.C. Jensen, S. Kloeker, C.L. Wang, M. Yoshigi, and M.C. Beckerle. 2006. Genetic ablation of zyxin causes Mena/VASP mislocalization, increased motility, and deficits in actin remodeling. J Cell Biol. 172:771–782.

Ibar, C., E. Kirichenko, B. Keepers, E. Enners, K. Fleisch, and K.D. Irvine. 2018. Tension-dependent regulation of mammalian Hippo signaling through LIMD1. J Cell Science. 131:jcs214700.

Jagannathan, R., G.V. Schimizzi, K. Zhang, A.J. Loza, N. Yabuta, H. Nojima, and G.D. Longmore. 2016. AJUBA LIM Proteins Limit Hippo Activity in Proliferating Cells by Sequestering the Hippo Core Kinase Complex in the Cytosol. Mol Cell Biol. 36:2526–2542.

Kim, N.-G., and B.M. Gumbiner. 2015. Adhesion to fibronectin regulates Hippo signaling via the FAK-Src-PI3K pathway. The Journal of Cell Biology. 210:503–515.

Kim, T.-J., S. Zheng, J. Sun, I. Muhamed, J. Wu, L. Lei, X. Kong, D.E. Leckband, and Y. Wang. 2015. Dynamic visualization of α-catenin reveals rapid, reversible conformation switching between tension states. Current biology. 25:218–224.

Krause, M., E.W. Dent, J.E. Bear, J.J. Loureiro, and F.B. Gertler. 2003. Ena/VASP Proteins: Regulators of the Actin Cytoskeleton and Cell Migration. Annual Review of Cell and Developmental Biology. 19:541–564.

Leerberg, J.M., G.A. Gomez, S. Verma, E.J. Moussa, S.K. Wu, R. Priya, B.D. Hoffman, C. Grashoff, M.A. Schwartz, and A.S. Yap. 2014. Tension-sensitive actin assembly supports contractility at the epithelial zonula adherens. Curr Biol. 24:1689–1699.

Li, J.X.H., V.W. Tang, and W.M. Brieher. 2020. Actin protrusions push at apical junctions to maintain E-cadherin adhesion. Proceedings of the National Academy of Sciences of the United States of America. 117:432–438.

Lin, V.T.G., and F.-T. Lin. 2011. TRIP6: An adaptor protein that regulates cell motility, antiapoptotic signaling and transcriptional activity. Cellular signalling. 23:1691–1697.

Marie, H., S.J. Pratt, M. Betson, H. Epple, J.T. Kittler, L. Meek, S.J. Moss, S. Troyanovsky, D. Attwell, G.D. Longmore, and V.M. Braga. 2003. The LIM protein Ajuba is recruited to cadherin-dependent cell junctions through an association with alpha-catenin. J Biol Chem. 278:1220–1228.

Misra, J.R., and K.D. Irvine. 2018. The Hippo Signaling Network and its Biological Functions. Annual Reviews of Genetics. 52:3.1–3.23.

Petit, M.M., S.M. Meulemans, and W.J. Van de Ven. 2003. The focal adhesion and nuclear targeting capacity of the LIM-containing lipoma-preferred partner (LPP) protein. J Biol Chem. 278:2157–2168.

Rauskolb, C., E. Cervantes, F. Madere, and K.D. Irvine. 2019. Organization and function of tension-dependent complexes at adherens junctions. Journal of Cell Science. 132:jcs224063.

Rauskolb, C., S. Sun, G. Sun, Y. Pan, and K.D. Irvine. 2014. Cytoskeletal tension inhibits Hippo signaling through an Ajuba-Warts complex. Cell. 158:143–156.

Razzell, W., M.E. Bustillo, and J.A. Zallen. 2018. The force-sensitive protein Ajuba regulates cell adhesion during epithelial morphogenesis. J Cell Biol:jcb.201801171.

Reinhard, M., K. Jouvenal, D. Tripier, and U. Walter. 1995. Identification, purification, and characterization of a zyxin-related protein that binds the focal adhesion and microfilament protein VASP (vasodilator-stimulated phosphoprotein). Proc Natl Acad Sci U S A. 92:7956–7960.

Rogge, H., N. Artelt, N. Endlich, and K. Endlich. 2017. Automated segmentation and quantification of actin stress fibres undergoing experimentally induced changes. J Microsc. 268:129–140.

Sanz-Rodriguez, F., M. Guerrero-Esteo, L.-M. Botella, D. Banville, C.P.H. Vary, and C. Bernabéu. 2004. Endoglin regulates cytoskeletal organization through binding to ZRP-1, a member of the Lim family of proteins. The Journal of biological chemistry. 279:32858–32868.

Schmidt, G., P. Sehr, M. Wilm, J. Selzer, M. Mann, and K. Aktories. 1997. Gln 63 of Rho is deamidated by Escherichia coli cytotoxic necrotizing factor-1. Nature. 387:725–729.

Smith, M.A., L.M. Hoffman, and M.C. Beckerle. 2014. LIM proteins in actin cytoskeleton mechanoresponse. Trends in cell biology.

Sun, S., and K.D. Irvine. 2016. Cellular Organization and Cytoskeletal Regulation of the Hippo Signaling Network. Trends in cell biology. 26:694–704.

Takizawa, N., T.C. Smith, T. Nebl, J.L. Crowley, S.J. Palmieri, L.M. Lifshitz, A.G. Ehrhardt, L.M. Hoffman, M.C. Beckerle, and E.J. Luna. 2006. Supervillin modulation of focal adhesions involving TRIP6/ZRP-1. J Cell Biol. 174:447–458.

Willier, S., E. Butt, G.H.S. Richter, S. Burdach, and T.G.P. Grunewald. 2011. Defining the role of TRIP6 in cell physiology and cancer. Biology of the cell. 103:573–591.

Yao, M., W. Qiu, R. Liu, A.K. Efremov, P. Cong, R. Seddiki, M. Payre, C.T. Lim, B. Ladoux, R.e.M.M.e. ge, and J. Yan. 2014. Force-dependent conformational switch of α-catenin controls vinculin binding. Nature communications. 5:4525.

Yi, J., and M.C. Beckerle. 1998. The human TRIP6 gene encodes a LIM domain protein and maps to chromosome 7q22, a region associated with tumorigenesis. Genomics. 49:314–316.

Yonemura, S., Y. Wada, T. Watanabe, A. Nagafuchi, and M. Shibata. 2010. alpha-Catenin as a tension transducer that induces adherens junction development. Nat. Cell Bio. 12:533–542.

Zanconato, F., M. Cordenonsi, and S. Piccolo. 2016. YAP/TAZ at the Roots of Cancer. Cancer cell. 29:783–803.

